# A Fast and Intuitive Method for Calculating Dynamic Network Reconfiguration and Node Flexibility

**DOI:** 10.1101/2022.02.06.479287

**Authors:** Narges Chinichian, Johann D. Kruschwitz, Pablo Reinhardt, Maximilian Palm, Sarah A. Wellan, Susanne Erk, Andreas Heinz, Henrik Walter, Ilya M. Veer

## Abstract

Dynamic interactions between brain regions, either during rest or performance of cognitive tasks, have been studied extensively using a wide variance of methods. Although some of these methods allow elegant mathematical interpretations of the data, they can easily become computationally expensive or difficult to interpret and compare between subjects or groups. Here, we propose an intuitive and computationally efficient method to measure dynamic reconfiguration of brain regions, also termed flexibility. Our flexibility measure is defined in relation to an a-priori set of biologically plausible brain modules (or networks) and does not rely on a stochastic data-driven module estimation, which, in turn, minimizes computational burden. The change of affiliation of brain regions over time with respect to these a-priori template modules is used as an indicator of brain network flexibility. We demonstrate that our proposed method yields highly similar patterns of whole-brain network reconfiguration (i.e., flexibility) during a working memory task as compared to a previous study that uses a data-driven, but computationally more expensive method. This result illustrates that the use of a fixed modular framework allows for valid, yet more efficient estimation of whole-brain flexibility, while the method additionally supports more fine-grained flexibility analyses restricted to biologically plausible brain networks.

## 1 Introduction

Over the past decades, a paradigm shift has taken place in studying the human brain, moving from a local to a more network-based perspective, giving rise to the field of network neuroscience. This evolution has, in part, been driven by the concept of graphs in math. A graph (network) consists of a set of vertices (nodes), which are connected by edges (links). In neuroimaging-based network neuroscience, brain regions identified by any given method of parcellation are considered the nodes of the network, while links can either be defined as white matter connections between brain regions (structural networks) or as statistical interdependencies between the time series of brain regions (functional networks) [6, 14, 17, 37, 38, 40, 41].

Mesoscopic structures or groups formed by interactions between nodes of a network, called modules, clusters or communities, can be quantified by a variety of detection methods [18]. Nodal interactions are typically represented by an adjacency matrix (A) of the network, where each element *i,j* of *A* (called *a_i,j_*) is the weight of the connection or strength of interaction between nodes i and j. Modules are usually determined based on the general idea of maximizing the number/weight of within-group and minimizing the number/weight of between-group links. Modules can then be considered as entities in the network that can be modified individually without affecting the rest of the network. Modularity measures have been shown to be useful as a biomarker of disease, including epilepsy [12], Alzheimer’s disease [9], schizophrenia, bipolar, and major depressive disorder [27]. However, brain modularity has also been associated with normal variation in cognition: Individuals with lower whole-brain modularity performed better in complex tasks, while those with higher modularity showed an advantage in simple tasks [46]. Whereas the ‘static’ community detection methods employed in the above-mentioned studies consider the brain’s connectivity averaged over time (based on only one adjacency matrix per subject as a single-layer network), other methods have assessed changes in community structure over time [1, 2, 10, 28, 32, 42]. These dynamic approaches take into account that a node can frequently change its connections depending on which state the brain is in, both during resting-state (RS) and during the performance of tasks. Here, changes in modular structure are captured by a sequence of adjacency matrices (*A_t_*), thus creating multi-layer networks. The adjacency matrices are typically calculated using a sliding-window approach on nodal time series, in which the window length reflects the time scale of interest [17]. Subsequently, dynamic module detection methods can be applied to these time-dependent multi-layer networks to not only characterize changes of modules over time, but also to determine how nodes change their affiliation [the module/group they belong to] as a function of time. The latter can be thought of as the flexibility of a node [2, 4] and is defined based on the consecutive presence of nodes in different modules over time [10, 29]. These measures of flexibility enable us to track time-dependent changes and thereby track phenomena of both integration and segregation in the brain [2, 7]. It offers the opportunity to study which brain nodes are more likely to change their affiliation over time and thereby which brain regions are rather consistently associated with a certain brain module, forming a backbone for the constantly changing network. For example, a recent study by Harlalka et al [21] suggested higher symptom severity in autism spectrum disorder to be associated with more connectivity flexibility in visual and sensorimotor areas during rest. Braun et al [7] demonstrated that individuals with more connectivity flexibility in frontal cortices have enhanced memory performance and score better on neuropsychological tests measuring cognitive flexibility, suggesting that dynamic network reconfiguration may form a fundamental mechanism underlying executive function. For a broader discussion on modularity and flexibility findings, see [24].

A data driven widely used method to calculate brain network flexibility is based on the Louvain community detection algorithm by Blondel et al [5]. This algorithm aims to optimize the variable *Q,* initially introduced for a single layer network in [33] by Newman, and later modified for multi-layer networks by others [3, 31, 44].

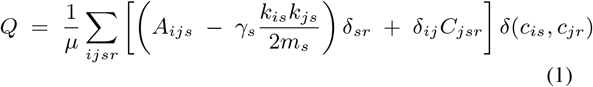

More specifically: Where *A* is the Adjacency matrix of the network, *A_ijs_* is the weight of connection between nodes *i* and *j* in layer *s. γ_s_* is the resolution parameter for layer *s, i* and *j* are indices of nodes, and *s* and *r* indices of layers. *k_is_* is the degree of node *i* in layer *s. m_s_* is proportional to the sum of weights in layer *s. C_jsr_* refers to the connection of node *j* to itself in different layers. *c_is_* is the defined module/cluster of node *i* in layer *s*. Finally, *Q* captures how good the grouping is compared to a null-model (here random).

Although, this and similar methods have undoubtedly contributed to our understanding of brain dynamics, these come with a cost: Given the random nature of algorithms like Louvain, the resulting clusters may differ each time the algorithm is run on the same adjacency matrix. As such, brain modules show variation within and across participants, which is overcome by running the algorithm multiple times to reach a consensus on the modular structure [25]. However, this can be a computationally expensive process, while the identified modules may in the end have low biological plausibility or at least can not be interpreted straightforwardly.

Here, we introduce a new method to capture nodal flexibility and brain network reconfiguration using a fast and intuitive method based on a set of template modules. This offers three main advantages over the existing methods:

1. It is computationally more efficient and deterministic compared to the Louvain (and similar) algorithm.
2. It offers high replicability, as it uses the same set of module templates for all subjects and time scales. This ensures comparability between subjects and studies, which is one of the current concerns in the field [20].
3. It gives researchers the opportunity to choose the best-fitting, or biologically most relevant module templates for each study.

In this work we describe our proposed method in detail and apply it to a real-life dataset that was previously assessed using a Louvain-like locally greedy heuristic algorithm [5, 7]. Compared to the previous work, we demonstrate that our method is equally successful in capturing a brain reconfiguration pattern that mimics the stimulation periods of an externally-cued working memory task, yet in our case can be directly related to well-known functional brain networks as well.

## 2 Method

### 2.1 Concept and Steps

Before going into mathematical detail, let us first explain) the concept behind the method. Consider the brain as a network, in which each region of the brain (defined by any arbitrary parcellation) is a node, each co-activation between any two nodes is an edge, and each node belongs to an a-priori defined set of nodes, termed a module. As a first step, we consider that each node has an a-priori affiliation to one of the predefined template modules or in other words, belongs to an a-prioiri template module. The affiliation is determined as the template module with which each node has the largest spatial overlap. Next, the strengths of all edges between each node and all members of every module are summed. When a node is more strongly connected to nodes affiliated with another module than to nodes of its own predefined module, then this node will receive another affiliation than its a-priori one. This can now be extended to a dynamic scenario, in which node affiliations can be determined for a range of consecutive time points. Some nodes might change their affiliation over time, while others do not. The ratio of nodes changing affiliation with respect to all nodes is what we are interested in. We understand this ratio as a measure of flexibility of the brain. In other words, the more nodes switch affiliation between consecutive time points, the more flexibility in network dynamics we assume. See Figure 1 for a summary of these steps.

**Figure 1:**
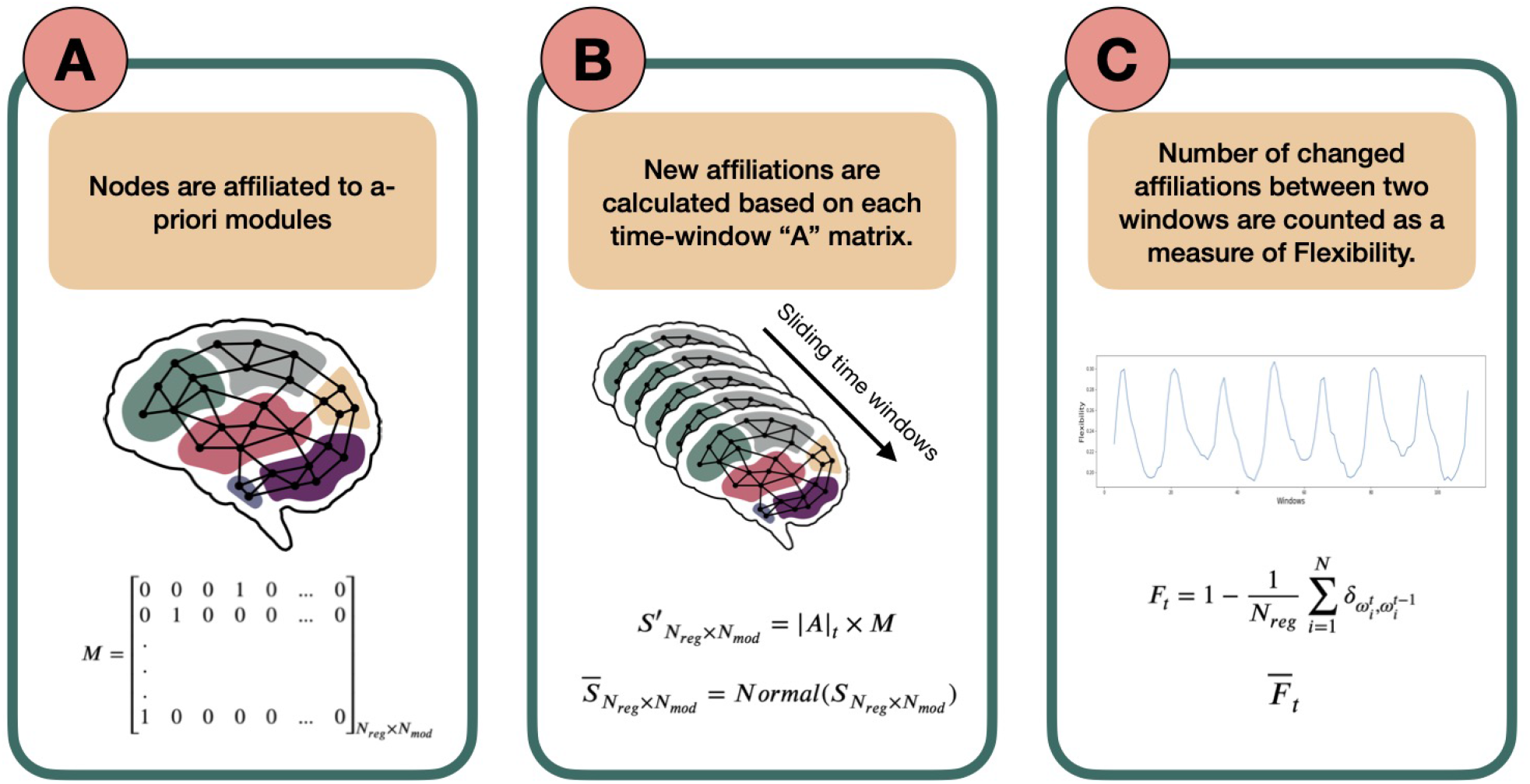
Schematic overview of the template-based flexibility method. A) Each node has an a-priori affiliation to a template module, not allowing overlap. In this paper, we use the Brainnetome atlas for node definition [15] and the FIND Lab network templates as predefined modules (http://findlab.stanford.edu/; [39]). Importantly, matrix M, describing the a-priori module affiliation for each node, is predetermined and serves as a reference. B) Using a sliding-window approach, an adjacency matrix is constructed for each time window by calculating Pearson correlation coefficients between the time series of all possible pairs of nodes. Then, for each node and time window the reference module receiving the highest normalized connection weight will serve as the new modular affiliation for that node in that time window. C) Last, the number of affiliation changes between affiliation vector in *t* and its successive vector in *t* + 1 is defined as the flexibility *F_t_* of the network between two time points. The average of *F_t_* across participants (called 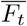) can be plotted for all consecutive time points (an example presented later in figure 3).

The steps to calculate this flexibility measure are listed below in detail:

1. An a-priori affiliation is assigned to each node to form the following matrix M:

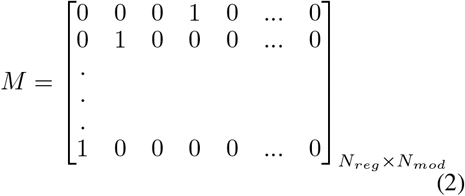 Where *N_reg_* is the number of regions (nodes) and *N_mod_* number of a-priori modules. Each row of this matrix belongs to a node and, in the first-approximation case in this paper, has only one non-zero element that indicates the a-priori modular affiliation of the node. For example, in row 1 the fourth column is 1, which means that the first node has an a-priori affiliation to template module 4. Note that we assign all nodes that do not show any overlap with the template modules to a last, artificial module to not exclude these nodes in calculating the flexibility metric.
2. Next, for each node we extract the mean time series across all volumes of the fMRI scan. We then divide our time-series into smaller windows using a sliding-window approach. For each time window, an adjacency matrix is constructed using Pearson correlation coefficients between all possible node pairs. The adjacency matrix at timewindow *t* is defined as *A_t_* of shape *N_reg_* × *N_reg_*:

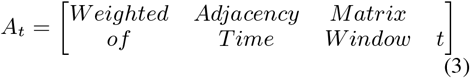
3. Now, we want to calculate how each node is connected to the nodes that are the predefined members of each of the template modules, as defined in *M*. To this end, we sum the absolute values of all the weights from one node to all the nodes affiliated to each of the modules, so that each node has *N_mod_* [in our subsection 2.2 analysis: 15] different values (one weighted sum for links to each module), indicating the strength of its links with the predefined members of each of the template modules. In mathematical terms, we calculate the matrix *S′* as follows:

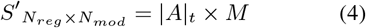

where |*A*|_*t*_ matrix elements are the absolute values of *A_t_* elements and the matrix has the dimension *N_reg_* × *N_reg_*. Row i of *S*′ belongs to the region *i* and each column *j* shows the sum of absolute connection weights of *i* to the members of *j*-th module. As the predefined modules differ in size, the *S*′ matrix elements are then normalized to the number of regions affiliated by template definition to the modules, creating a new matrix called *S* [dividing each matrix element 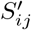 by the number of regions affiliated to the *j*th template module.] Importantly, to be able to compare the elements of *S*, we normalize it in a way that the sum of each row is one. This normalization step has no effect on the output of the next steps but is rather to increase the interpretability at this stage. The normalized numbers thus represent which portion of each node’s connections is to which module. We call this new matrix, 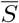.

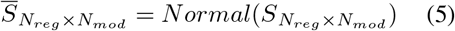
4. With 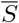, we have the ratio of affiliations to each module calculated for all nodes. From these, the strongest module affiliation per node is chosen as the winner which together form an affiliation vector for time window t; we call this vector Ω_*t*_:

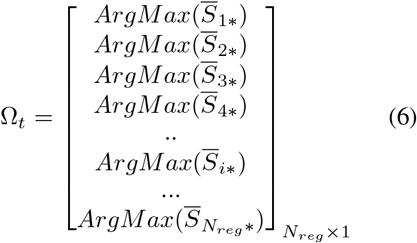

where 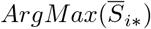 points to the name/number (argument) of the winner module in row i of matrix 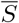.
5. Following steps 2-4 for consecutive time windows, we calculate one Ω_*t*_ for each window t. The *flexibility* of the network denoted by *F* is then defined as the ratio of regions that change their affiliation from one window to the next to the total number of network regions, or:

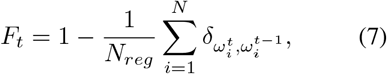

where *ω* denotes an element of vector Ω. The Kronecker delta 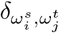 is 1 if 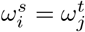 and 0 otherwise. The then counts the number of nodes that did not change their affiliation between windows *t* and *t* + 1. Note that as a side-product of calculating Ω, we can output a vector describing the affiliations over time for each node separately as well by making a vector of the same element in Ω_*t*=1_,..,*N_t_*:

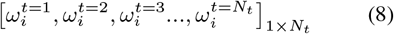

Where *N_t_* is the total number of time windows. This output can be used for further region-specific analysis.
6. Where we apply the method to real-life data (see subsection 2.2) we also calculate the average *flexibility* over time for a sample (cohort of subjects), 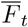, by simply summing the flexibility over all participants and divide it by the sample size (*N_sub_*).

### 2.2 Application on a Previously Studied Dataset

In our application study, we used 331 participants of the 344 participants included in [7]: Thirteen subjects were excluded due to scanning artefacts, exceeding movement or insufficient image quality. Functional MRI data were acquired at three sites during performance of an N-back task: the Life and Brain Center of the University of Bonn, the Central Institute of Mental Health Mannheim, and Charité - Universitätsmedizin Berlin. The study was approved by the Medical Ethics Committees of the three study sites and all participants provided written informed consent. At all sites, a Siemens Trio 3T MRI scanner (Siemens Healthcare, Erlangen, Germany) was used with identical sequences: gradient-echo EPI, 28 slices, slice thickness 4mm (1mm gap), field of view 192 x 192 x 140mm, acquisition matrix 64 x 64, TR (repetition time) 2s, TE (echo time) 30ms, flip angle 80°. The task was presented in a blocked fashion. Four blocks of 0-back and 2-back each (30s duration) were alternated, starting with the 0-back condition. Participants were asked to either press the button corresponding to the number shown on the screen (0-back) or the number that was shown 2 steps ago (2-back). See Figure 2 for more information on the task. Standard preprocessing was conducted using SPM8 [36] and included motion correction (participants with > 3mm translation and > 1.7° rotation between volumes were excluded), slice-time correction, spatial smoothing with a FWHM of 9mm, high-pass temporal filtering with a 128s cutoff, and normalization to the Montreal Neurological Institute (MNI) template space with 3mm isotropic voxel size. A detailed description of data acquisition and preprocessing is provided in [13]. Mean time-courses of the 246 Brainnetome Atlas regions [15] were extracted from the preprocessed data of the 331 subjects. In line with [7], a 15-volume window length with 14 volumes overlap was chosen for the sliding-window analysis (figure 2.C and 2.D), generating in total 114 windows for each subject. For every window, we calculated an adjacency matrix using Pearson correlation coefficients between all possible pairs of the 246 regions mean time series (using scipy.stats.pearsonr [45]). Considering that the N-back working memory task consisted of 30s alternating blocks of 0-back and 2-back, the 15-volume window (30s length) allows for one window purely reflecting a single condition block. For more information on selection of the window length see [7] and [26].

**Figure 2:**
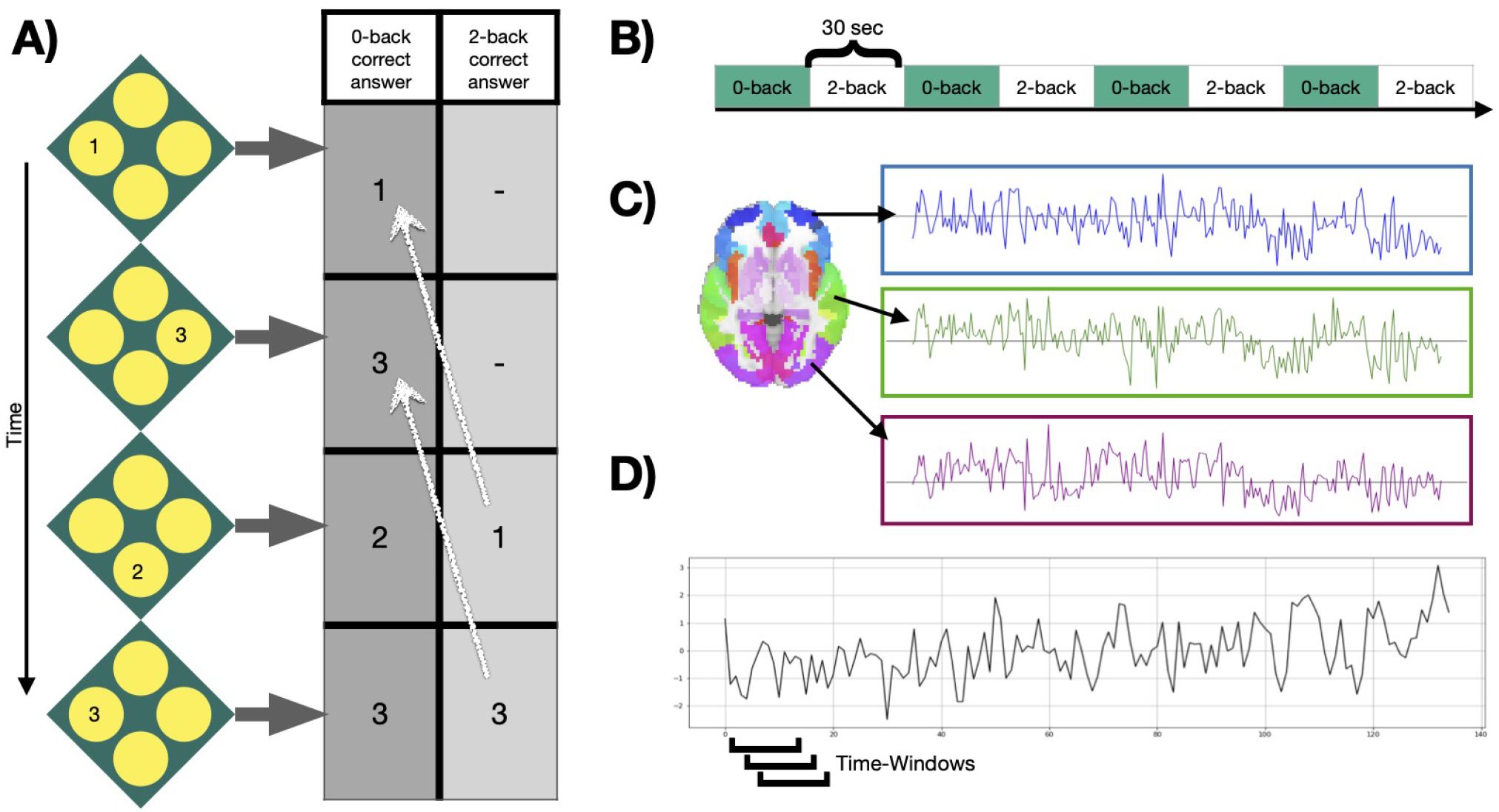
Task and signals. A) Example of the N-back working memory task with a 0-back and 2-back condition, during which participants were asked to choose the value that was either shown at the current step or 2 steps ago, respectively. B) Four blocks of each condition were presented in alternated fashion for 30 seconds. C) After preprocessing, mean time courses were extracted from 246 Brainnetome atlas regions [15]. D) Windowed time series were extracted using a sliding-window approach, moving a window of 15 time points over the time series one volume at a time.

The a-priori modules (Matrix M) were selected based on 14 well-described functional connectivity template networks (modules) in [39] by the FIND lab (http://findlab.stanford.edu/). As described before, a 15th (artificial) module was added comprising all atlas regions that did not overlap with any of the 14 template networks. The a-priori affiliations of all atlas regions can be found in table 1 and the labels of the FIND lab templates in table 2.

**Table 1:**
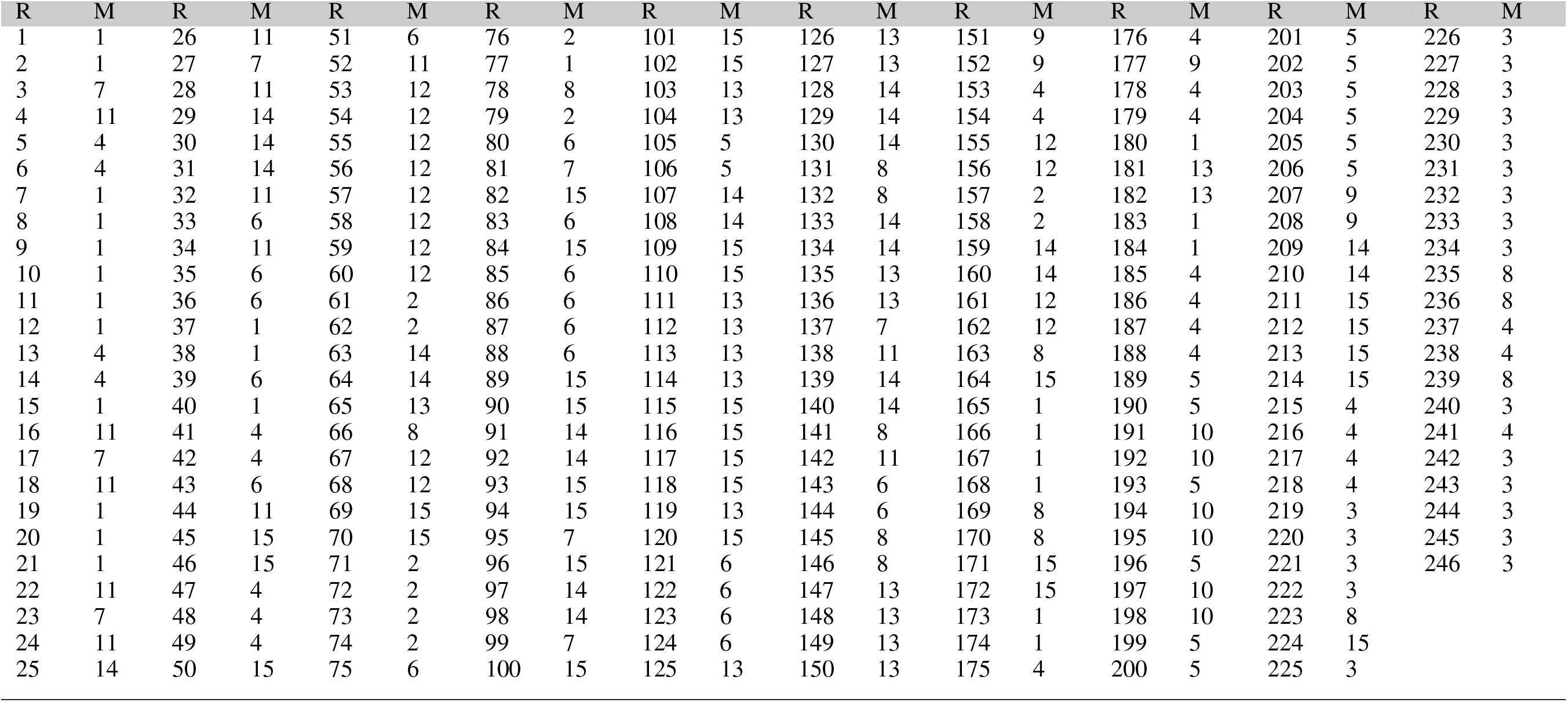
Region a-priori Affiliation, columns marked “R” are region numbers and “M” columns are a-priori modular affiliations

**Table 2:** Findlab-based modules [39] used in our application section.

| Number | Name |
| --- | --- |
| Module 1 | Anterior Saliency |
| Module 2 | Auditory |
| Module 3 | Basal Ganglia |
| Module 4 | Dorsal Default Mode Network (dDMN) |
| Module 5 | High Visual |
| Module 6 | Language |
| Module 7 | Left Executive Control (LECN) |
| Module 8 | Posterior Saliency |
| Module 9 | Precuneus |
| Module 10 | Prim Visual |
| Module 11 | Right Executive Control (RECN) |
| Module 12 | Sensorimotor |
| Module 13 | Ventral Default Mode Network (vDMN) |
| Module 14 | Task Positive |
| Module 15 | Undefined |

To obtain a broader view of the meso-scale dynamics, the modular allegiance matrix T and integration matrix R were calculated using the methods from [7]. Each element *t_i,j_* of modular allegiance matrix T shows the ratio of windows where node i and j were present in the same module relative to all windows. To calculate the T for each condition, we separated windows with 80% of their time-points in one condition and ignored the others.

To calculate the integration matrix R with elements *r_k,l_,* which show the strength of co-working between modules k and l, when we have *N_mod_* modules {*M*_1_, *M*_2_,…*M_N_mod__*}, we first use all the T matrix elements [link between two regions] with one end (region) in module k and the other end (region) in module l to extract I matrix elements (*i_k,l_*). It can be written as:

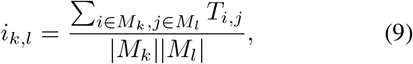

where k and l are two modules, |*M_k_*| shows the size of module *M_k_*. Then we normalize the I elements with division by internal connections of both modules and call the resulting elements elements of matrix R:

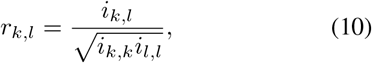

## 3 Results

Figure 3.A shows the N-back flexibility pattern across all nodes from [7], while Figure 3.B shows the pattern generated by our method when applied to the same dataset (331/344 subjects of the same sample). Similar to Figure 3.A, the peaks illustrate maximum flexibility of the brain during performance of both the 0- and 2-back condition. In contrast, the transitions between the two task conditions coincide with troughs when applying our method, whereas [7] described additional, yet smaller peaks during these transition phases when using the generalized Louvain algorithm. On average, higher flexibility is observed during the 2-back than 0-back blocks, although the difference is relatively small (*t* = −2.9, *p* = .03).

**Figure 3:**
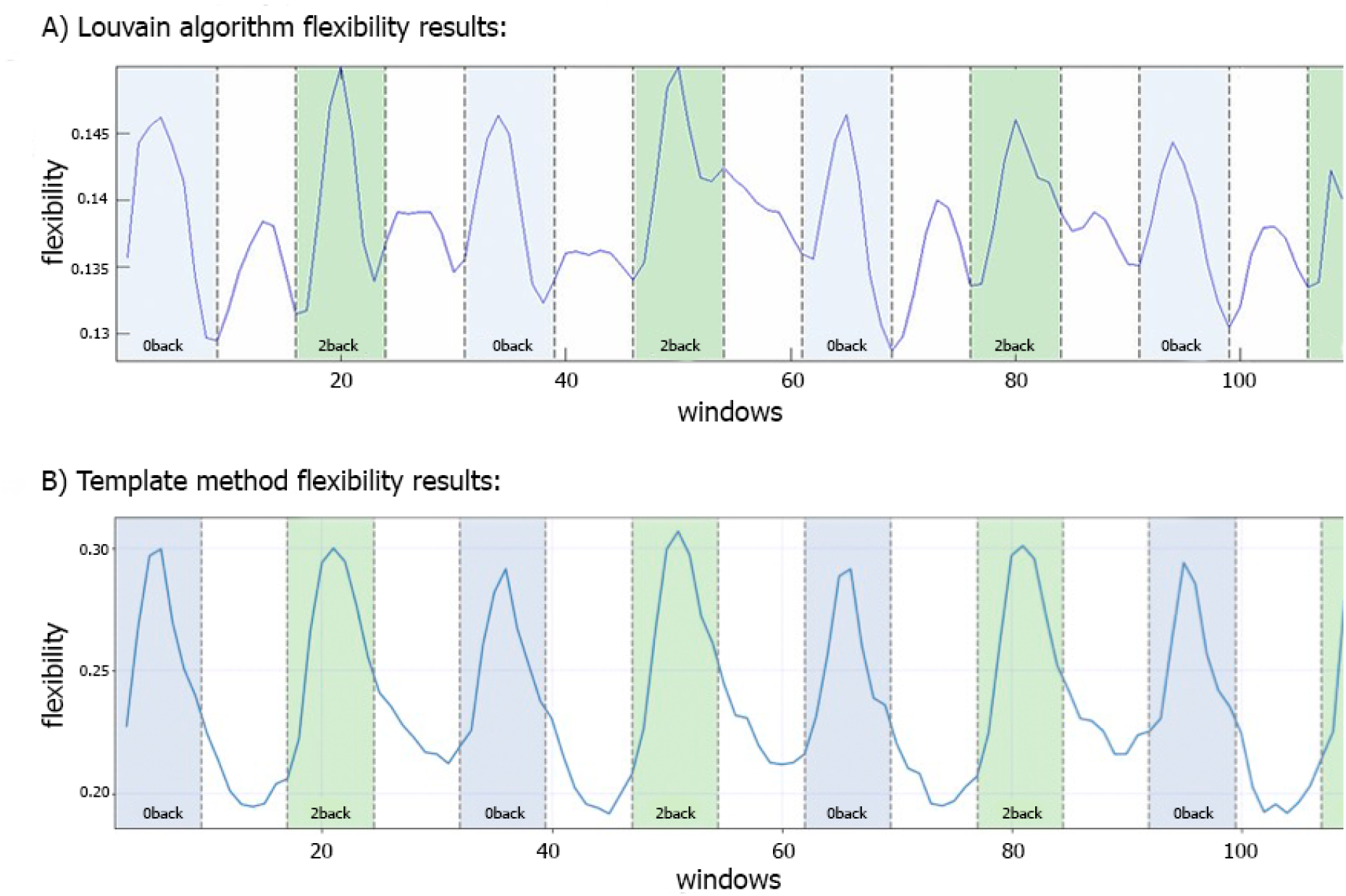
Comparison of flexibility generated by the generalized Louvain-like locally greedy heuristic algorithm [5,23] and the template-based method during an N-back working memory task. A) Flexibility plot from [7] illustrating the probability that a brain region changes its modular allegiance between two consecutive windows in a sample of 344 healthy subjects. The original plot is used with permission of the publisher. B) Flexibility plot generated by the template-based method. Here, the flexibility number in each time-window is the fraction of regions that change their affiliation from one time window to the next (i.e., the number of changed regions divided by the total number of nodes). The plots are generated using a subset of 331 subjects from the same cohort as used in [7]. Note that in both plots a time window covers 15 EPI volumes with a TR of 2s, corresponding to a window length of 30s. The window was shifted with one volume at a time, allowing for 14 EPI volumes overlap between consecutive windows, which yielded 114 windows in total.

In addition to calculating flexibility across all nodes, we can use the information captured in the fifth step to describe the affiliation changes of each individual node. This allows us to have a closer look at which nodes switch their affiliation over time most frequently, or at how often the a-priori constituents of each of the template networks switch their affiliation. Figure 4 illustrates how many times each node (Brainnetome regions in our analysis) switches its affiliation between two consecutive windows. Note that the number of switches was normalized to the number of switches performed by the node that switched most frequently, forcing the latter node to have a value of 1 and the other nodes to have a value between 0 and 1. Nodes within the prefrontal cortex predominantly show affiliation changes over time during execution of the N-back task. This is in agreement with the previous findings [8, 11, 30, 34].

**Figure 4:**
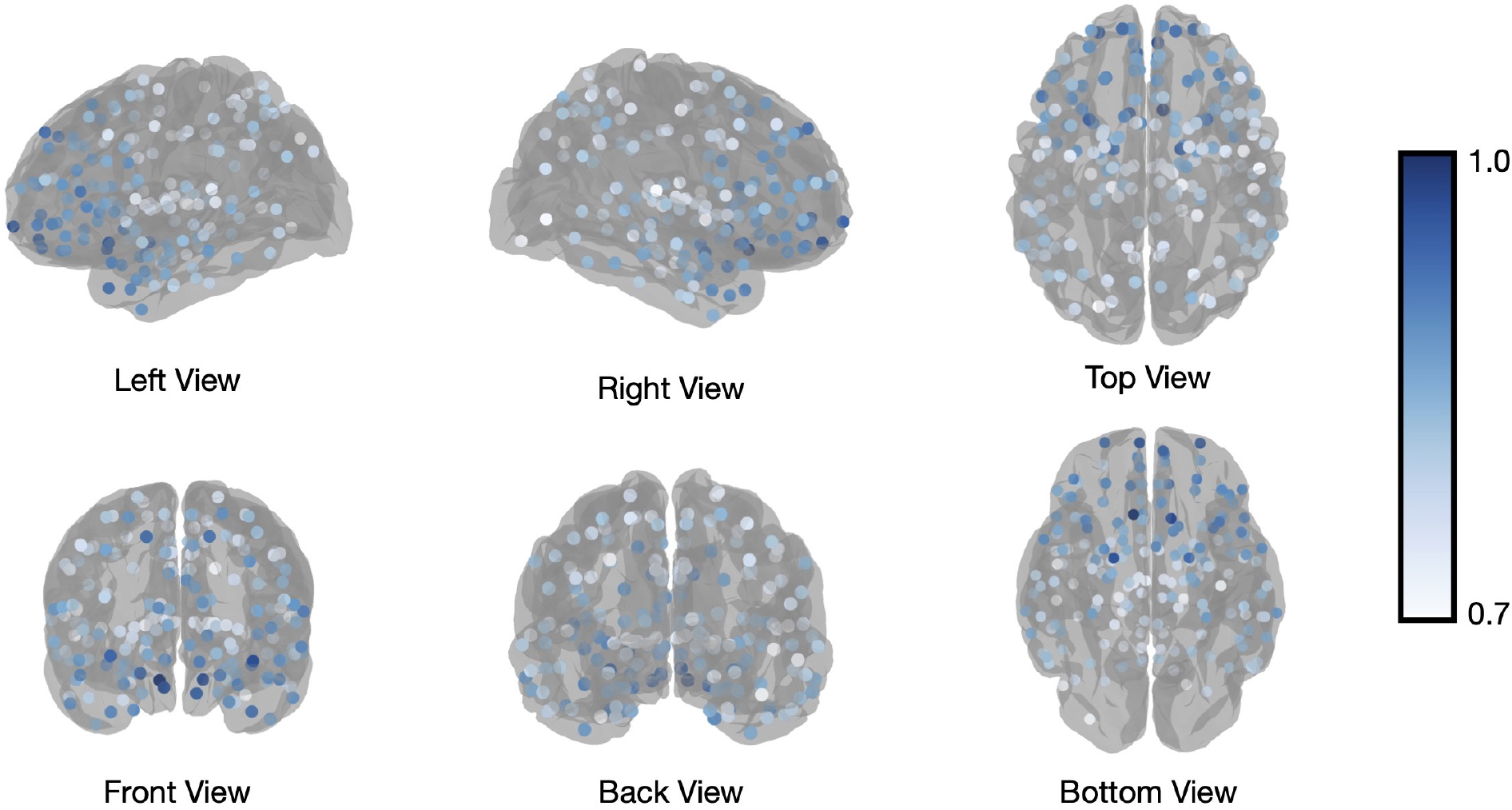
Brannetome atlas brain regions switching. Number of affiliation switches between consecutive windows for regions of the Brainnetome Atlas, averaged across all subjects and normalized to the most frequently switching node to yield values between 0 and 1. The visualized regions are those with values higher that 0.7.

One level coarser at the module level, we can look at the average switching ratio of template modules. The boxplots in Figure 5 demonstrate for each of the FIND lab template modules how often their a-priori defined constituent nodes on average switch their modular affiliation over time across participants. Constituent nodes of the default mode network (DMN), salience network (SN), left and right executive control network (L/RECN), and language network seemingly switch their affiliation most often during execution of the N-back task.

**Figure 5:**
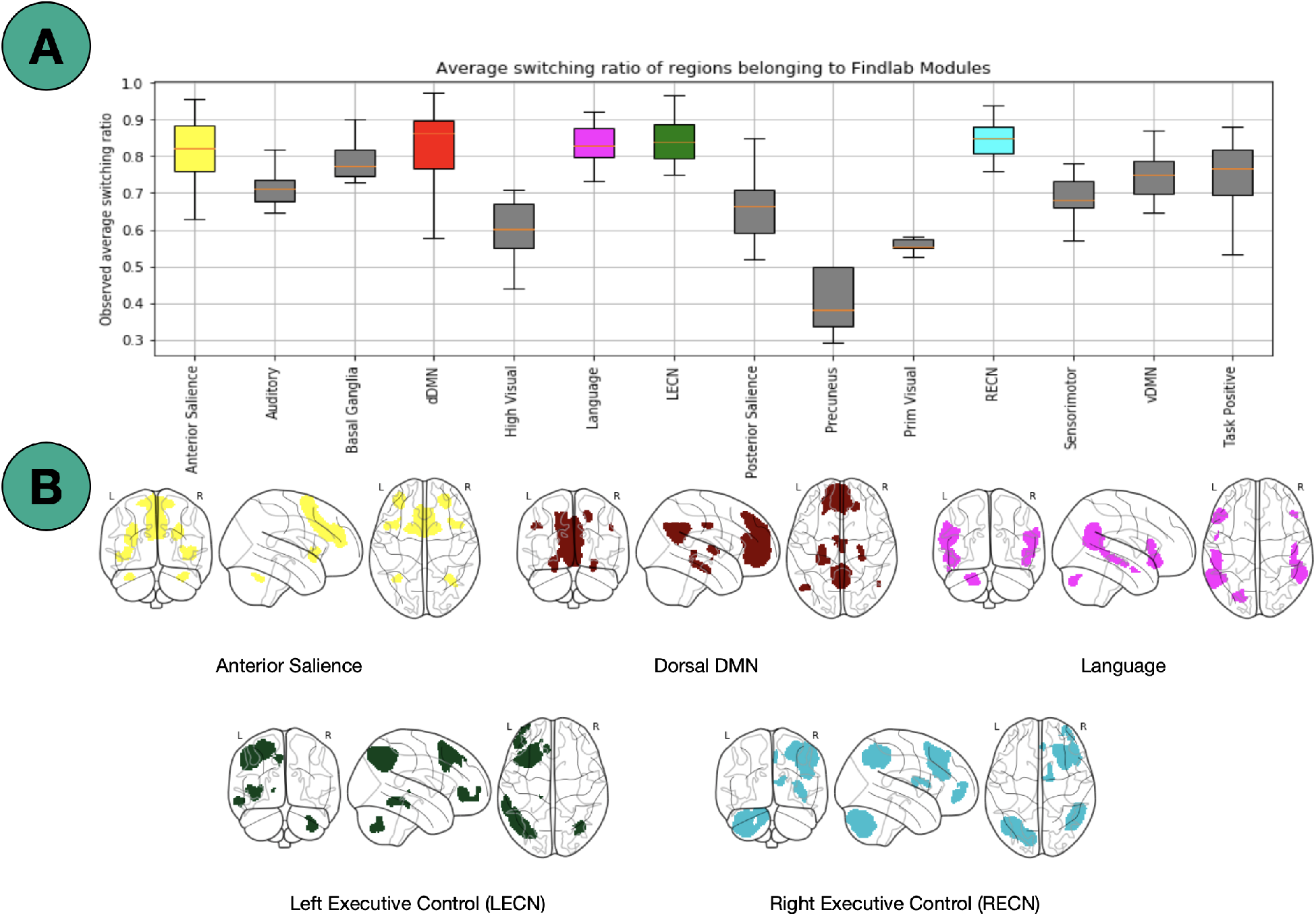
Findlab brain areas switching. A) Average number of affiliation switches between consecutive windows for each FIND lab template network, averaged across all subjects. Abbreviations are listed in table 2. B) Illustration of the four template networks for which its constituent nodes demonstrated the highest flexibility (http://findlab.stanford.edu/; [39]).

Figure 6 shows the result of modular allegiance and integration analysis. We observe a general increase in integration values in 2-back compared to 0-back except for 3 modules. This overall increase in integration is in agreement with previous findings [16].

**Figure 6:**
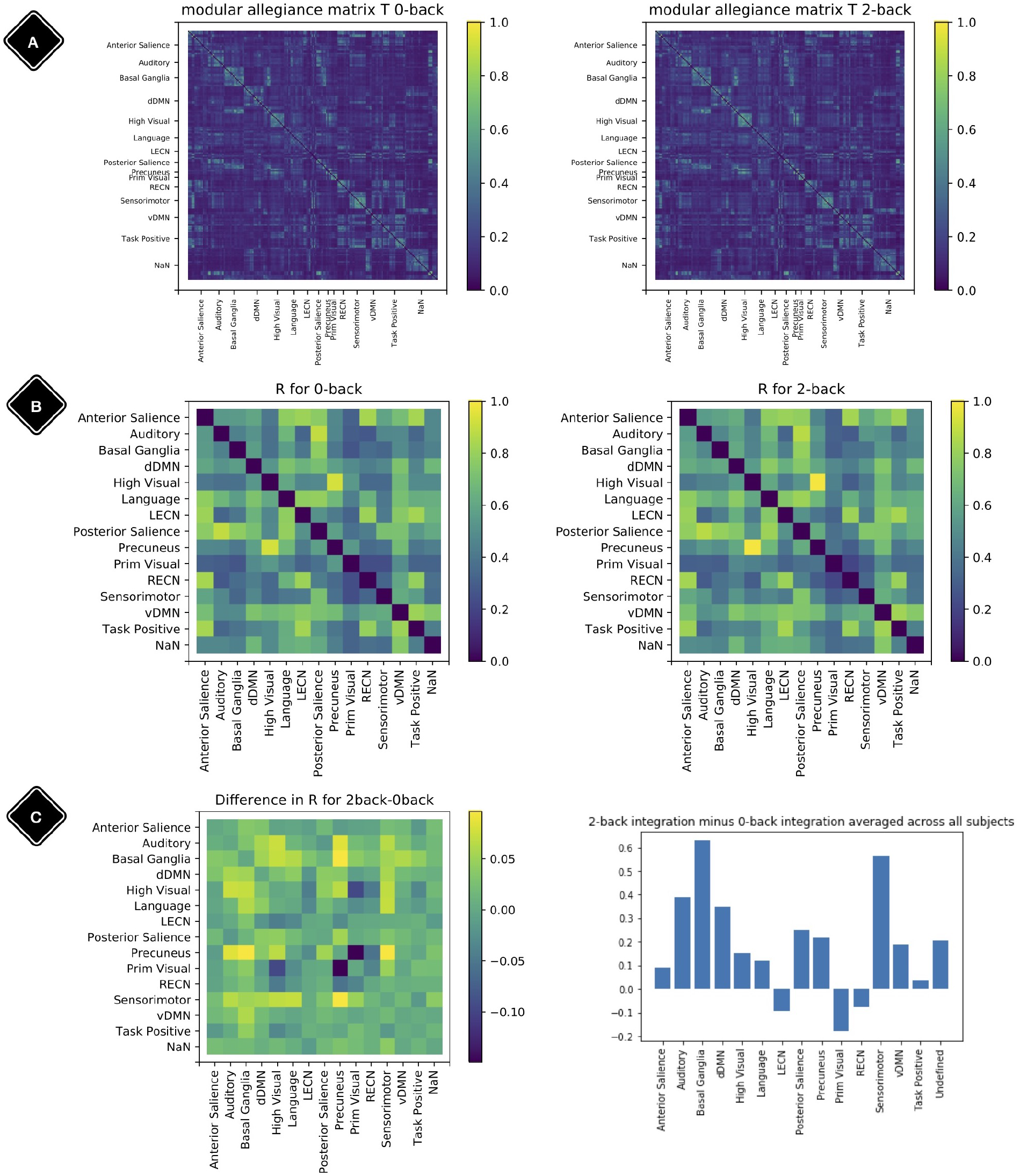
Modular allegiance and integration. Diagonal elements of the matrices are set to be zero. A) Modular allegiance of the two conditions 2-back and 0-back; to calculate a T matrix for one condition, we used only the windows with 80% of their time-points in that condition. B) Integration matrix for 0-back and 2-back. C) Change in the integration values *R*_2-*back*_ – *R*_0–*back*_ (left plot) and sum of rows (from the left plot matrix) as each modules integration value (right plot).

## 4 Discussion

In this work we introduce a new method to assess flexibility in analyses of dynamic functional connectivity. In the application section we set out to compare our method against the currently most used data-driven method described in [7], in which the computationally more expensive generalized Louvain algorithm was applied to derive the modular structure of the data [2, 5, 23, 31]. We demonstrate that our method is able to reveal a flexibility pattern during the N-back working memory task that is highly similar to the pattern found in [7]. The most notable difference between the results obtained with our method and the Louvain algorithm was the absence of the small increase in flexibility during the transition of the 0- and 2-back blocks. Braun and colleagues [7] interpret this to reflect “dual-task” performance. We suggest an alternative explanation based on the current results: increased flexibility may be needed for switching tasks at the start of each new condition block (shown as a delayed peak in the middle of the marked blocks), while less flexibility may be needed during prolonged execution of the task in each block (shown as a delayed trough exactly in between blocks). As such, the periods of lower flexibility may show the preferred brain configuration for the execution of the task blocks. A further more theoretical analysis of a simulated BOLD signal with block induced inputs might be helpful in interpreting the dual-task vs. no-dual-task hypothesis.

As has been shown abundantly in the literature, the prefrontal cortex plays an important role in the performance of working-memory tasks [8, 11, 30, 34]. Therefore, it is not surprising that we found nodes in the prefrontal cortex to show the most flexible behavior during execution of the N-back task. Moreover, at the modular level we see the highest flexibility in nodes that have an a-prori affiliation to the DMN, SN, L/RECN and language modules. The DMN is known to have an antagonistic relation with frontoparietal networks, such as the L/RECN: when the latter is more active during cognitively demanding tasks (such as the N-back) the DMN is less active [19]. Interestingly, a key role has been assigned to the SN in allocating neural resources between more internally (DMN) or externally (ECN) oriented processes [43]. Taken together, we see these results as further proof of our method’s validity.

We discussed above how our method could be used to assess flexibility. That is, both on the network (module) level and at the regional (node) level, thereby extending the inferential potential compared to the other widely-used algorithms. However, our analytical procedure also offers possibilities for more fine-grained investigations of modular affiliations. In the description of our method and application analysis we determined the modular affiliation for a particular node and window as the module with which the node demonstrated the strongest connectivity in the affiliation vector. Although this is arguably the easiest and most pragmatic choice, it would also be possible to use the weighted affiliation with each of the template modules in the affiliation vector [Method section, step 3] to assess flexibility. Such a weighted approach may ultimately prove to be even more informative in characterizing brain flexibility.

In conclusion, the method proposed in the current study is able to generate flexibility results that are highly comparable to the results obtained with a more sophisticated data-driven method. Besides having a much higher computational efficiency, our method also promotes replicability across different samples and studies through the use of biologically plausible template modules. We believe that our approach can be a feasible choice for researchers aiming to study dynamical reconfiguration at multiple scales of the brain, be it nodes, modules, or the brain as a whole.

## A Acknowledgements

This study was supported by the Deutsche Forschungsgemeinschaft (DFG, German Research Foundation) - SPP2041, WA 1539/9-1 /SPP2031, WA 1539/11-1, ERK 724/4-1.

We thank Professors Eckehard Schöll and Tilo Schwalger from TU Berlin together with their group members for their constructive contributions to improve this method and manuscript.

## B Visualization

Python packages “nilearn”, “scikit-learn” and “Matplotlib” are used for the visualizations [22, 35].

## C Codes

The Python scripts for our method are available on GitHub: https://github.com/NargesChinichian/FlexDraftPub Please contact the first author on GitHub or via email for further questions.

